# Neurite Density and Free Water in the Gray and White Matter of Early Psychosis Patients

**DOI:** 10.1101/2024.09.09.612136

**Authors:** Peter C. Van Dyken, Ali R. Khan, Lena Palaniyappan

## Abstract

Diffusion weighted imaging has been frequently used to characterize the white matter in patients with schizophrenia, but the most commonly used model, diffusion tensor imaging (DTI), is not specific to the histological nature of microstructural changes. This is particularly true in the more complex grey matter tissue. Furthermore, DTI changes have not been consistently reported in early schizophrenia populations, but this does not exclude more subtle changes that may not affect the model fit. Recently developed biophysical models of diffusion, such as the neurite orientation dispersion and density imaging (NODDI) model, may overcome these issues by quantifying specific tissue subcompartments, capturing diffusion profiles characteristic of intra-neurite, extra-neurite, and free water space. We applied the NODDI model to early schizophrenia patients (n=54) and healthy controls (n=51) from the Human Connectome Project - Early Psychosis dataset, investigating both the grey and white matter. We observed a diffuse, increased free water fraction throughout the grey matter, especially in the left insula, though there were not notable changes in the white matter. The spatial variation in the grey matter free water was not fully explained by the partial volume effects from the cerebrospinal fluid, indicating a role for tissue edema. The role of vasogenic processes in early stages of psychosis that may precede white matter anomalies documented in later disease stages warrant further investigation.

## 1 Introduction

Chronic schizophrenia has been robustly associated with altered diffusion tensor imaging (DTI) parameters in the white matter, but this disruption is not consistently present in the first 5 years since diagnosis (1–14). Thus, DTI alterations may not be a phenotype of schizophrenia as such, but more related to chronic forms of the disease.

DTI parameters, however, are a very nonspecific measure of white matter pathology, reducing a sizable volume of tissue into a single three-dimensional ellipsoid. Several factors can thus account for DTI changes, and some processes may not cause any changes at all. For instance, in regions with prominent crossing tracts, fractional anisotropy (FA) will be low *a priori* in healthy controls (HCs). In such regions, disrupted tract integrity in a disease may not result in FA changes.

This creates three problems for the interpretation of DTI parameters in schizophrenia. First, the underlying pathophysiological process causing DTI changes cannot be directly determined. Second, it is not clear if subtle white matter changes, such that do not affect DTI, are present in early schizophrenia. Third, DTI is not suitable for application to the grey matter, with its much more complex tissue architecture.

These problems may, in part, be resolved by newer biophysical models of diffusion. These capitalize on higher resolution imaging to make more specific statements about tissue microstructure (15). Of these, neurite orientation dispersion and density imaging (NODDI) has become quite popular, both for its rich information and because its acquisition protocol is within reach for most major imaging centres (16). NODDI decomposes the diffusion signal into three additive compartments (Figure 1): a free-water, isotropic compartment (*v_iso_*), an extra-cellular, anisotropic compartment, and an intracellular, stick-like compartment. Both the intra- and extra-neurite compartments account for directionality, making the model robust to crossing fibres. Using these compartments, composite metrics may be computed including neurite density index (NDI), the intra-cellular compartment normalized by the total constrained diffusion, and orientation dispersion index (ODI), which measure the directional homogeneity of the intra-cellular signal.

**Figure 1:**
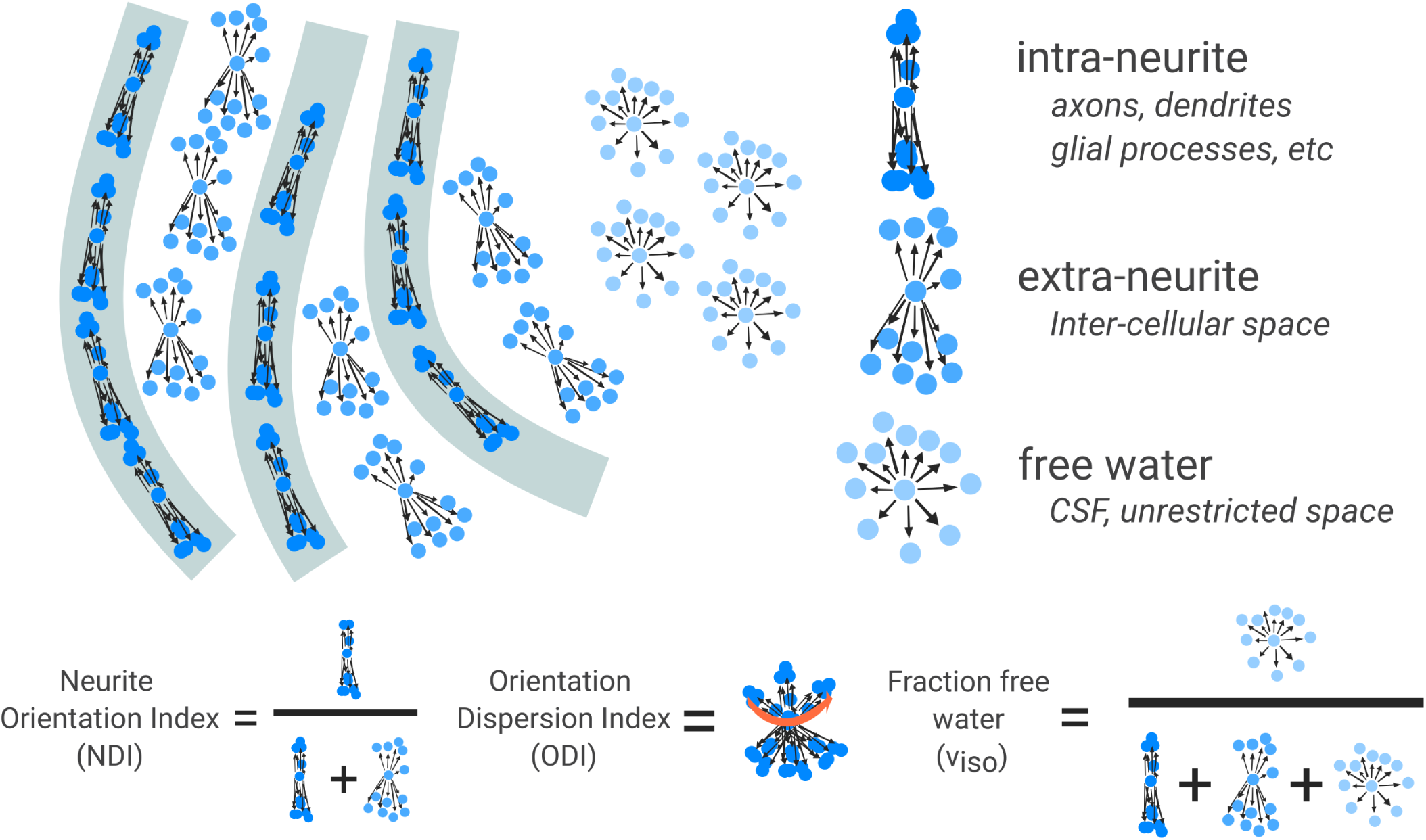
NODDI models three tissue compartments that, together, compose the observed diffusion signal. An intra-neurite compartment is composed of stick-like diffusion signal, an extra-neurite compartment is ovoid diffusion signal, and free water is spheroid signal. These compartments are fit to the emperical signal and assigned a weight. From these weights, summary measures can be computed. Neurite orientation index (NDI) is the amount of intra-neurite signal as a proportion of all neurite-derived signal (excluding free water). Orientation dispersion index (ODI) indexes the directional heterogeneity of intra-neurite (stick-like) diffusion within the voxel. Low ODI corresponds to directionally coherent intra-neurite signal, while high ODI corresponds to multiple, directionally diverse components. Fraction free water (*v_iso_*) is the weight of free water signal as a proportion of signal from all compartments. This represents water that diffuses without barriers, such as CSF.

Because of its compartmental specificity, NODDI is an attractive choice for modelling the grey matter, in addition to white matter analysis. Changes in NDI or *v_iso_* can be more readily interpreted in terms of neuropil and ganglionic density.

NODDI is still relatively new, and although its acquisition requirements are much simpler than other biophysical models, it is still more involved than the imaging protocols used in many clinical studies. It has thus not yet been extensively used in imaging studies of schizophrenia (17). Studies in chronic patients have reported reduced NDI in the dorsolateral prefrontal cortex and temporal lobe (18,19) with heterogeneous ODI changes (19,20). *v_iso_* increases and decreases were varyingly observed in white matter tracts (18). Two studies have looked at NODDI in the white matter of first episode psychosis patients. Their results are contrasting, with one reporting NDI reductions in patients with no ODI changes (21), and the other showing mixed NDI increases and decreases with widespread ODI increases (22). No study to our knowledge has investigated the grey matter.

This report presents findings from an analysis of the HCP-EP dataset (23) using the NODDI model. Previous investigations of this dataset observed correlations between DTI metrics and negative symptoms of schizophrenia (1,24), despite no group-level differences in patients compared to controls (1). We hypothesize that the NODDI parameters — NDI, ODI, and *v_iso_* — will be more sensitive markers of schizophrenia and show extensive differences from controls even at an early stage of illness, both in the grey matter and white matter. We also hypothesize similar associations with negative symptoms will be observed. As a reference, we also investigate cortical thickness, a more extensively characterized marker that is consistently reduced in schizophrenia populations (25,26).

## 2 Methods

### 2.1 Data

Human Connectome Project - Early Psychosis (HCP-EP) data was accessed according to the Data Use Certification issued by the NIMH Data Archive. The patient group included subjects with DSM-5 diagnosis of schizophrenia, schizophreniform, schizoaffective, psychosis not otherwise specified, delusional disorder, or brief psychotic disorder within five years of initial diagnosis. Subjects with affective psychosis were excluded from this study. All diagnoses were made after six months of care. Healthy controls had no personal and family history of psychosis/schizophrenia, and subjects with a current, treated anxiety disorder were additionally excluded. Complete inclusion and exclusion criteria is retrievable (at time of publication) at https://humanconnectome.org/study/human-connectome-project-for-early-psychosis/ document/hcp-ep. Data was acquired on a 3-T MRI (Siemens MAGNETOM Prisma). T1-weighted (T1w) data was collected using an MPRAGE sequence at 0.8mm isotropic resolution, echo time = 2.22ms, repetition time = 2.4 s, field of view = 256 mm, number of slices = 208. Diffusion magnetic resonance imaging was acquired with an EPI sequence at 1.5mm isotropic resolution, echo time = 89.2 ms, repetition time = 3.23 s, field of view = 210 mm, number of slices = 92, MB acceleration factor = 2, flip angle = 78. 92 directions were acquired in both the AP and PA directions at b=1500 and b=3000, along with seven b=0 images.

### 2.2 Preprocessing

#### 2.2.1 Anatomical data

The minimally preprocessed anatomical images included in the dataset (27) were used for anatomical analysis.

#### 2.2.2 Diffusion Data

Diffusion data was preprocessed using *snakedwi* (28), a preprocessing pipeline based on *snakebids* (29) and *snakemake* (30). Briefly, Gibbs ringing artefacts were removed with *mrdegibbs* from *MRtrix3* (31,32); eddy currents and motion were corrected using *eddy* from *FSL* (33); susceptibility-induced distortions were corrected using *topup* in *FSL*, using the AP-PA pairs of images (34,35). The T1w image was skull stripped with *SynthStrip* (36); bias field correction was applied with *N4ITK* from *ANTS* (37). A T1w proxy image was created from the diffusion image using *SynthSR* (38) and used to accurately register the diffusion image space to the T1w space using a rigid transform calculated with *greedy* (39). DTI metrics were calculated using *dtifit* from *FSL*with linear regression (40), using all b-values for the fit.

Tractography was performed using the *MRtrix3* (32) software suite. Constrained spherical deconvolution was performed using the dhollander algorithm to estimate the response functions for white matter, grey matter and cerebrospinal fluid (CSF) (41,42). Multi-tissue constrained spherical deconvolution, as implemented in *MRTrix*, was performed to obtain white matter-like fibre orientation distributions (FODs) as well as grey matter-like and CSF-like compartments in all voxels (43). *mtnormalise* was used to correct for residual intensity inhomogeneities (44,45). Tractography was performed using the iFOD2 algorithm (46) and anatomically constrained tractography (ACT) (47), with 10,000,000 streamlines (48), an FOD amplitude cut-off of 0.06, a minimum streamline length of 4mm, and a maximum streamline length of 250mm. The anatomical segmentation used for ACT was obtained using *SynthSeg* (49). Spherical-deconvolution informed filtering of tractograms was used to correct the streamline counts based on the underlying FOD magnitude (50).

Subject anatomical scans were parcellated using the Brainnetome atlas (51). These parcellations were used to derive weighted connectivity matrices based on the tractography data. Average NDI, ODI, and *v_iso_* were used as weights.

### 2.3 NODDI

The NODDI model was fit using the *Microstructure Diffusion Toolbox* (52,53) with preprocessed images in the original acquisition space. The log *v_iso_* image was subsequently computed using the *v_iso_* map. Voxels equal to zero were eliminated by creating a smoothed version of the image, then assigning zero-valued voxels from the original image with their non-zero counterparts in the smoothed image. Other voxels were left unmodified.

NODDI parameter maps were transformed to the T1w image space using the rigid transform computed in *snakedwi*.

### 2.4 Surface Sampling

FA, mean diffusivity (MD), radial diffusivity (RD), axial diffusivity, NDI, ODI, and *v_iso_* were sampled from the grey matter and projected onto the *FSlR* cortical surface using the *-volume-to-surface-mapping* command from *wb_command*. We performed ribbon-constrained sampling between the pial and white matter surfaces, projecting onto the mid-thickness surface. Voxels were weighted based on their distance from the mid-thickness surface according to a Gaussian with sigma equal to 10% of the cortical thickness at the point of sampling. Additionally, to help limit partial voluming, a mask was created from the *v_iso_* map selecting all voxels with *v_iso_* < 0.1. Only voxels selected by this mask were sampled, for all parameter maps.

### 2.5 Skeletonization

FA maps were first non-linearly registered to a common template corresponding to the average space of all FA images. This template was computed with an in-house implementation of the iterative algorithm described by Avants *et al.* (54), using *greedy* (39) to compute each transformation. NODDI derived parameter maps (NDI, ODI, log *v_iso_*) were transformed into this space. *FSL* (35) was then used to skeletonize the maps.

### 2.6 Parcellations

Global surface averages were computed by averaging across all vertices for each parameter map, per hemisphere. The Desikan-Killiany atlas computed by *fastsurfer* was used to parcellate the surface data. Hemispheres were kept separate throughout the analysis.

A composite, hierarchical atlas was constructed using the Johns Hopkins University (JHU) atlas distributed with *FSL* (55) and the Talairach lobe segmentation (56). Both atlases were first intersected with the skeletonized image, and the lobe segmentation was used to define a peripheral atlas by subtracting the JHU atlas. Mathematically:

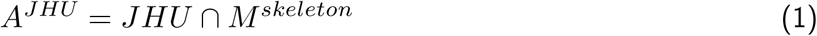

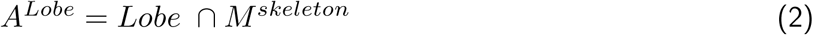

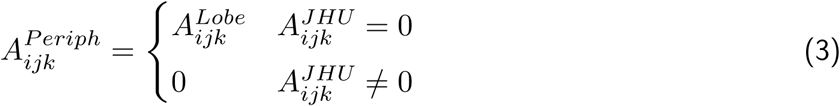

where *A_ijk_* is the *ijk*th voxel of the indicated atlas. Homologous ROIs across the two hemispheres were merged. This approach enabled us to recover the regions outside of the white matter core (defined by the JHU atlas) to define the periphery.

Four hierarchical levels were defined for statistical purposes:

1. The global mean (one ROI)
2. Mean of all voxels in the peripheral atlas and of all voxels in the JHU atlas (two ROIs)
3. Peripheral atlas and JHU atlas merged into four tract groupings (eleven ROIs)
4. JHU atlas (23 ROIs)

The tract groupings used in level 3 for the JHU atlas are shown in Table 1.

**Table 1:**
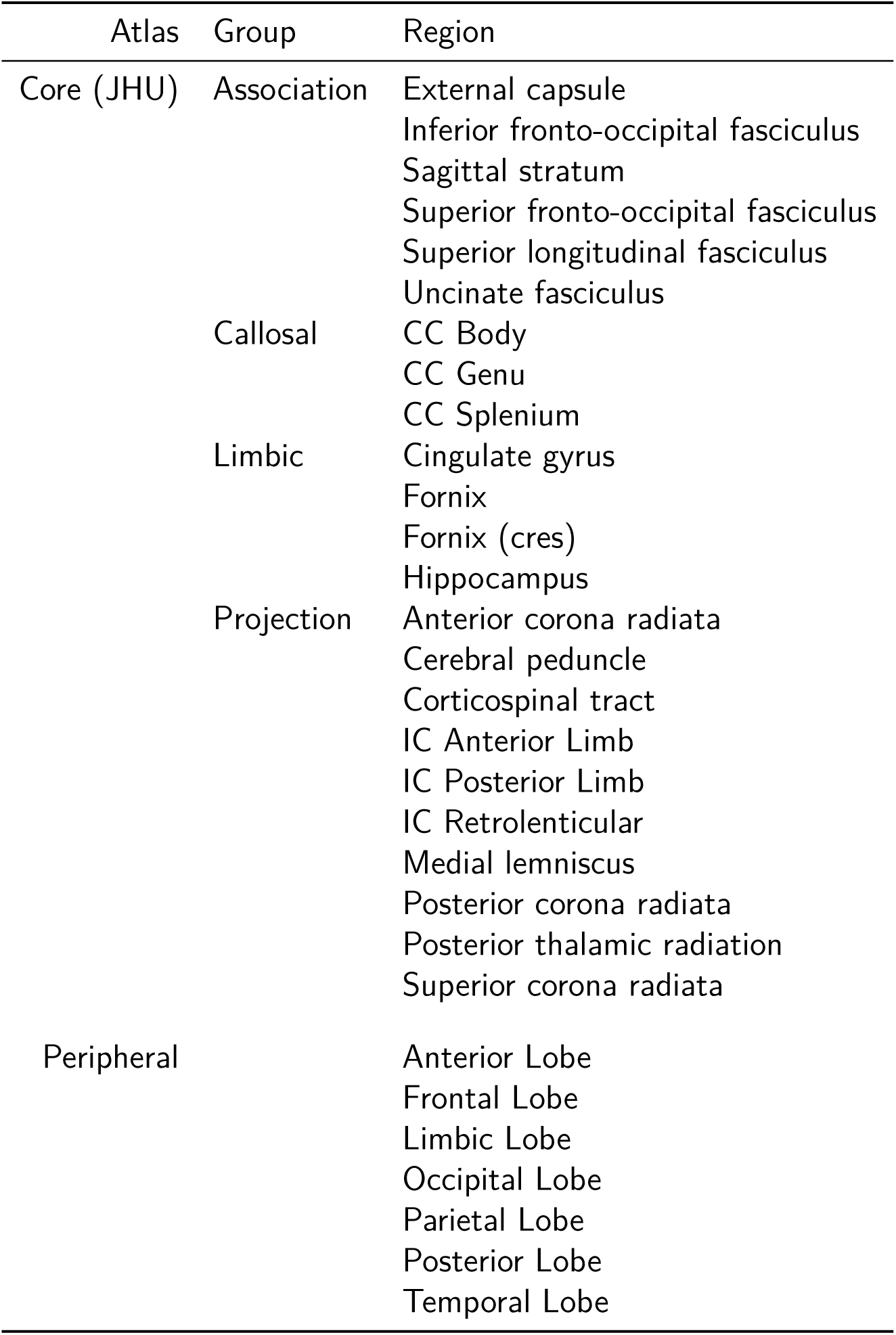
ROIs used in study, with the four subgroupings of the JHU atlas.

### 2.7 Analysis

Outliers were excluded as follows: for each metric (n=9), for each subject, the z-score for each ROI (n=7) was calculated relative to values from the same ROI, the same metric, and within the same group (patient or control) of subjects. The absolute value of these z-scores were then rounded down to the nearest integer. The rounded z-scores were summed across all ROIs and metrics for each subject. Subjects whose summed z-scores exceeded 30 were excluded from further analysis. This threshold was chosen qualitatively with the aim of excluding subjects with numerous metric-ROI combinations at a z-score of >3 while limiting operator bias.

White matter and grey matter ROIs were compared using two-sample T-tests, with age and sex as nuisance variables. For NDI, ODI, and thickness, one-way comparisons were performed testing for lower values in patient data. For *v_iso_*, one-way comparisons were performed testing for higher values. For DTI variables tested in the cortex, two-way comparisons were used.

Gray matter surface ROI comparisons were corrected for multiple comparisons using the Benjamini-Hochberg false discovery rate (FDR) procedure (57). For white matter ROIs, FDR correction was performed separately for each hierarchical atlas level, as previously performed in (58).

*BrainStat* was used to perform surface-based clustering of NDI and DTI parameters using random field theory (59–66). One and two-way comparisons were performed as described above. Age and sex were treated as nuisance variables. The cluster threshold for statistical significance was 0.01, family-wise error rate (FWER) was set as 0.05.

*BrainSpace* was used to perform spin tests (67) across parameter maps. For each permutation, spatial correlation was computed using Spearman rank-order correlation. A null distribution was computed using 1000 permutations, with two-tailed comparisons for significance testing.

Connectivity differences were assessed using the subject-specific connectomes. Edges with significantly differing weights across groups were identified using network based statistic (NBS) (68) with extent based cluster sizes, a T threshold of 3.0, 10,000 iterations, and corrected p-value threshold of 0.05. Sex and age (and site for HCP-EP) were regressed as nuisance variables.

Correlations between clinical variables and parameters were completed as described above, replacing the diagnostic group variable with the clinical variable of interest (and analysis was restricted to patients). We considered PANSS30-P, PANSS30-N, and a Disorganization score, computed from PANSS30 subscores as:

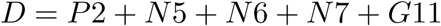

Disorganization is one of the most inconsistently defined factors in the PANSS score (69) with different items per analysis and distributed loading of core items. We chose the most consistent items (P2, N5, N7 and G11), dropping items that were infrequent in the sample (G10 Disorientation, G13 Disturbed volition). The co-occurrence of impoverished speech (N6) with disorganized speech, constituting formal thought disorder syndrome, has been observed in longitudinal studies in the early stages of psychosis (70) and with cluster analysis (71). N6 has therefore been included with other items of disorganization in the present study.

### 2.8 Code Availability

Unless otherwise indicated, all analyses and figures were generated with Python 3.11 using openly available python packages (72–82), particularly *pybids* (83,84), *nibabel* (85), and *nilearn* (86). Analysis code for this project can be obtained at https://github.com/pvandyken/study-hcp-noddi.

## 3 Results

### 3.1 Demographics

Three HCs and two patients were excluded as outliers (Section 2.7). One additional patient with unusable data was also excluded.

Patients were matched to HCs for sex and socioeconomic status but were were significantly younger (*t* = 4.18, *P* < 0.001) and more right-handed (*χ*^2^ = 3.45, *P* < 0.05). As expected in a population with schizophrenia, patients were less educated (*t* = 4.8, *P* < 0.05) and more likely to smoke (*χ*^2^ = 2.54, *P* < 0.05) than healthy counterparts. Cannabis use was not reported amongst HCs, but 20.3% of patients had at least some exposure. These patients were in treatment for less than 5 years, with variable medication compliance over this period (Table 2).

**Table 2:** HCP-EP demographics.

|  | HC (n=51) | Patient (n=54) | HC vs Patient |
| --- | --- | --- | --- |
| Sex (M/F) | 35/16 | 40/14 | $\chi^2(1) = 0.161, P = .69$ |
| Education | 15.71 (1.71) | 13.29 (1.31) | $T(36) = 4.8, P < .001$ |
| Age | 24.67 (4.06) | 21.81 (2.87) | $T(103) = 4.18, P < .001$ |
| Handedness (R/L) | 40/10 | 49/3 | $\chi^2(1) = 3.45, P = .063$ |
| SES | 2.10 (1.05) | 2.53 (1.23) | $T(101) = -1.89, P = .062$ |
| Smoke (Yes/No) | 3/46 | 10/44 | $\chi^2(1) = 2.54, P = .11$ |
| Pack years | 0.02 (0.10) | 0.29 (0.73) | $T(77) = -2.22, P = .029$ |
| Cannabis User (Yes/No) |  | 11/43 |  |
| Antipsychotic Duration (months) |  | 17.76 (15.65) |  |
| Lifetime Antipsychotics (Defined Daily Dose) |  | 223.15 (244.32) |  |
| PANSS Total |  | 51.83 (9.67) |  |
| PANSS Positive |  | 11.67 (3.77) |  |
| PANSS Negative |  | 15.55 (5.17) |  |
| PANSS Global |  | 24.75 (4.57) |  |

### 3.2 Grey matter

Globally, cortical thickness was significantly lower in patients, but no other group effects were found (group effects of lower NDI and higher *v_iso_* in patients did not survive Holm-Bonferonni corrections) (Figure 2 A). We observed significantly lower NDI in the right hemisphere across all participants (Figure 2 B), but no other hemispheric effects or group by hemisphere interactions (Figure 3). Effect sizes and T-values are given in Table 3. Because of the NDI differences, the hemispheres were considered separately in further analyses.

**Figure 2:**
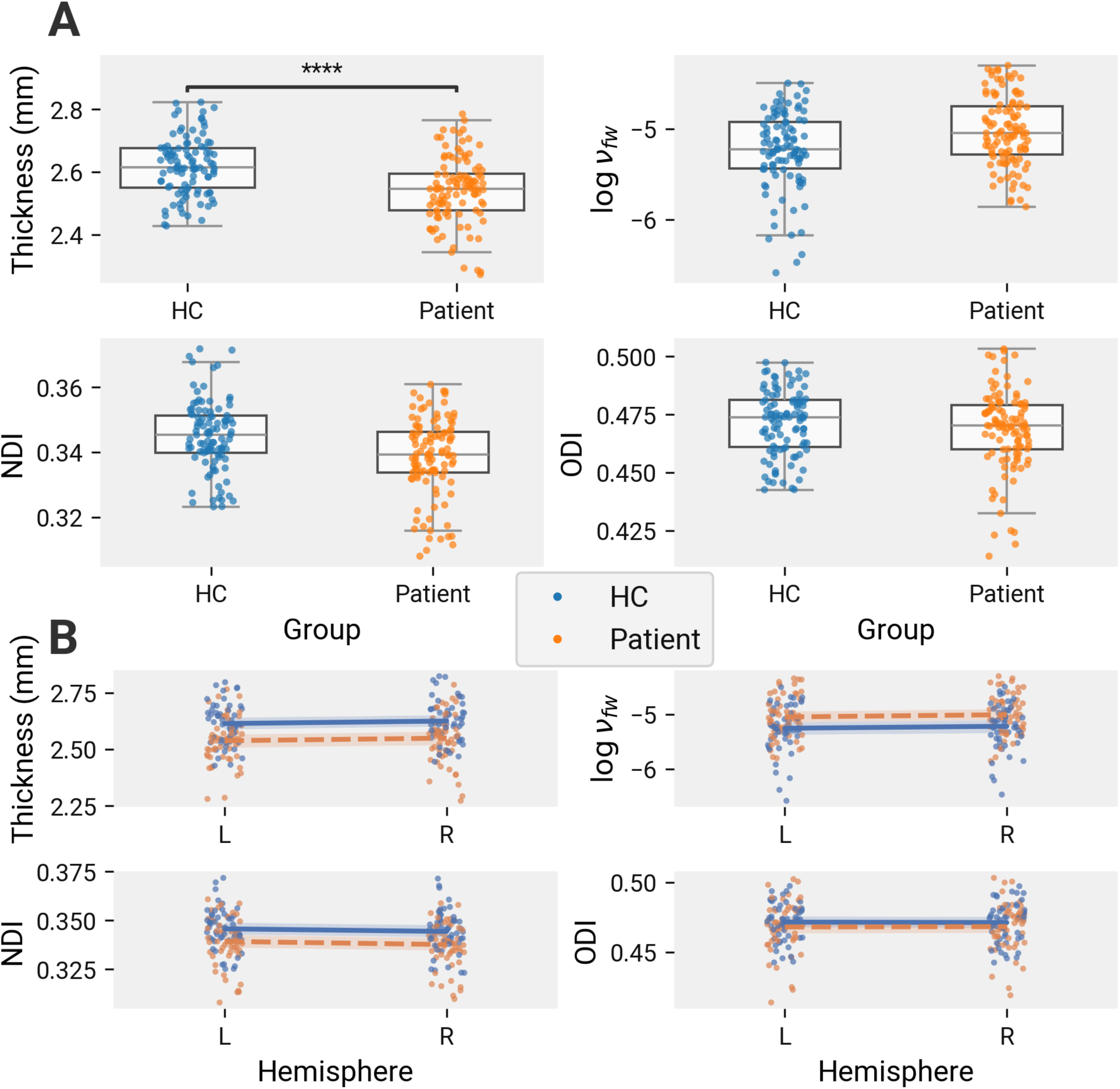
Cortical thickness is reduced in patients. Statistics computed using linear mixed-effects models with subject as a random effect and group, hemisphere, age, and sex as fixed effects. For each parameter, two-tailed T-tests were used to evaluate the effect of hemisphere and group-hemisphere interaction on the parameter. One-tailed T-tests were used for group effects on NDI, ODI, thickness (reduced in patients), and *v_iso_* (increased in patients). Degrees of freedom for each comparison was estimated using Satterthwaite’s method (87). P-values from the three contrasts were corrected using Holm-Bonferonni corrections. A. Average cortical thickness is significantly lower (*T* (110.4) = −4.8; *P* < .001) in patients versus controls. B. Average parameter values split by hemisphere. Lines illustrate the per-group mean differences across hemispheres. Shaded bands represent 95% CI computed by parametric boostrapping of the mixed-effects model with 1000 replicates. NDI is significantly lower in the right hemisphere (*T* (103.0) = 3.5; *P* = .002). No other hemisphere or group:hemisphere interaction effects are noted.

**Figure 3:**
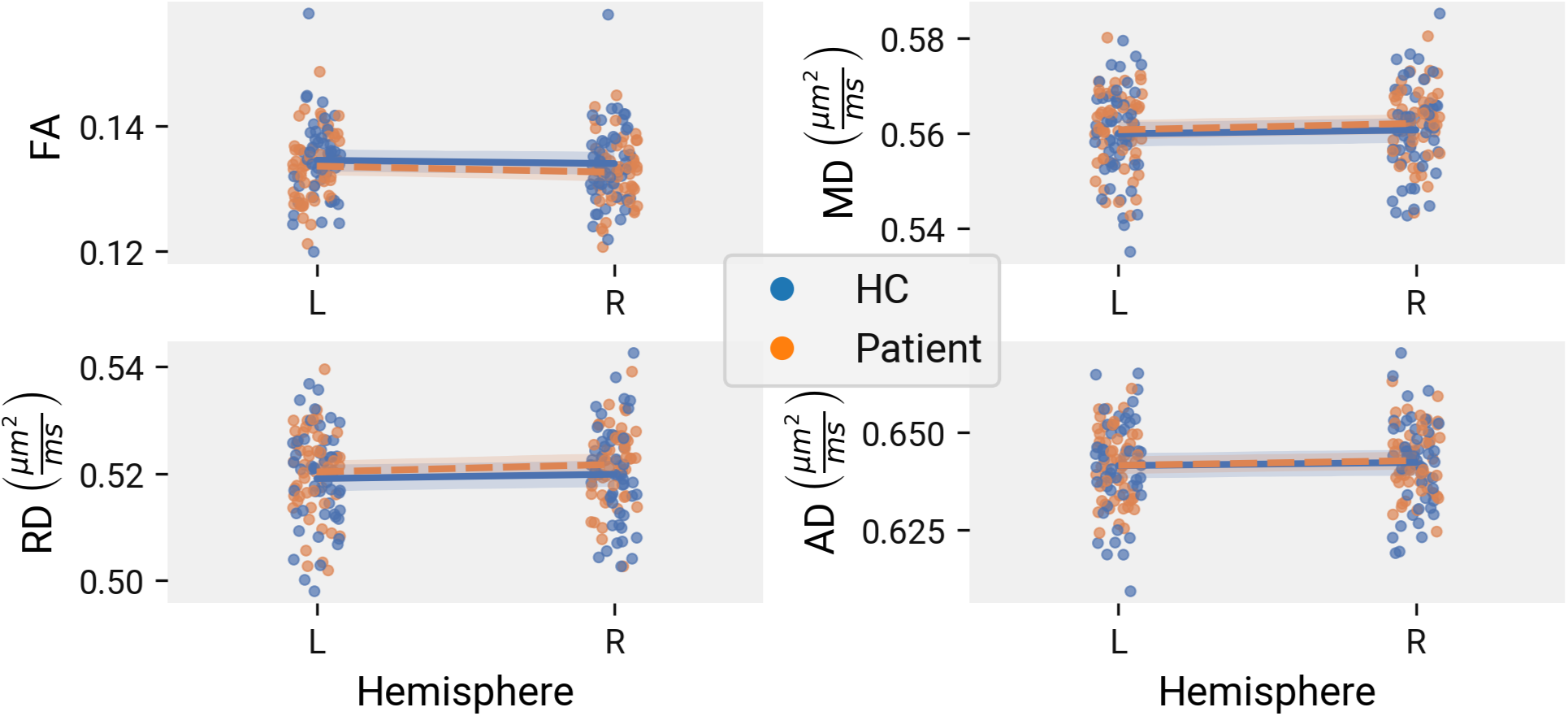
No DTI changes in patient cortical gray matter. Statistics computed using linear mixed-effects models with subject as a random effect and group, hemisphere, age, and sex as fixed effects. For each parameter, two-tailed T-tests were used to evaluate the effect of group, hemisphere, and group-hemisphere interaction on the parameter. Degrees of freedom for each comparison was estimated using Satterthwaite’s method (87). P-values from the three contrasts were corrected using Holm-Bonferonni corrections. Average parameter values are represented split by hemisphere. Lines illustrate the per-group mean differences across hemispheres. Shaded bands represent 95% CI computed by parametric boostrapping of the mixed-effects model with 1000 replicates.No hemisphere or group:hemisphere interaction effects are noted.

**Table 3:**
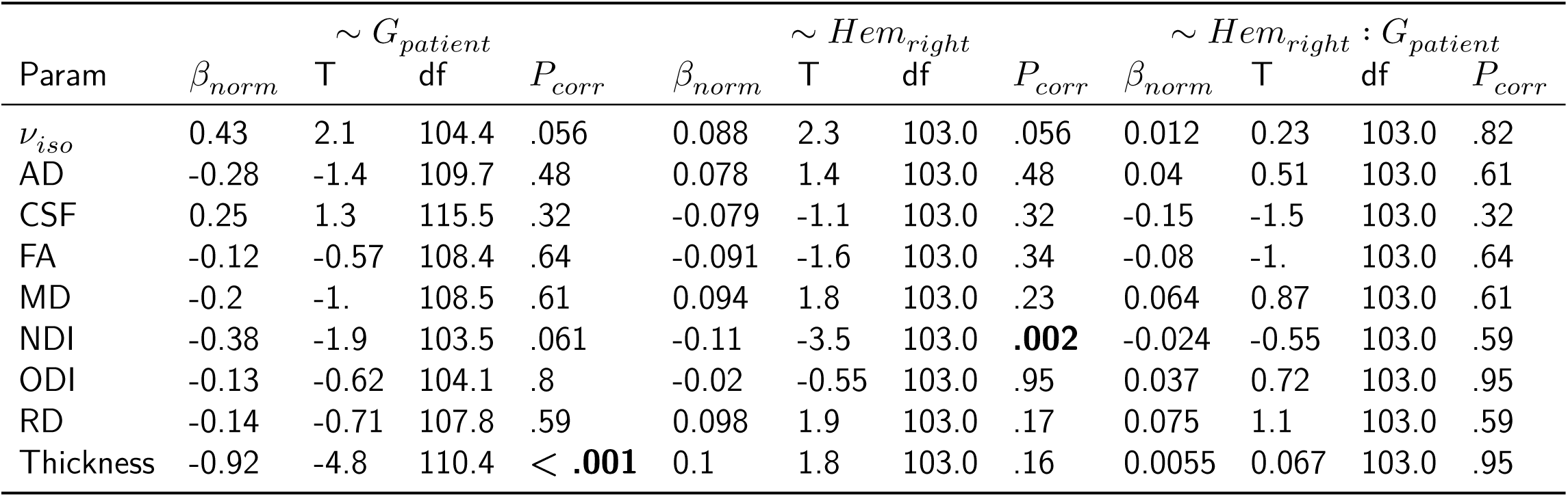
Statistics for average cortical parameters modelled against group and hemisphere.

Breaking down the hemispheres into ROIs, reduced thickness in patients was found in every cortical ROI in both hemispheres. Most ROIs had increased *v_iso_*, including the bilateral frontal, temporal, and occipital lobes, bilateral insulae, and the right cingulate gyrus (Figure 4 A). We additionally observed reduced FA in the left insula (Figure 4 B). No differences were found for NDI, ODI, or any of the other DTI parameters (Table 4).

**Figure 4:**
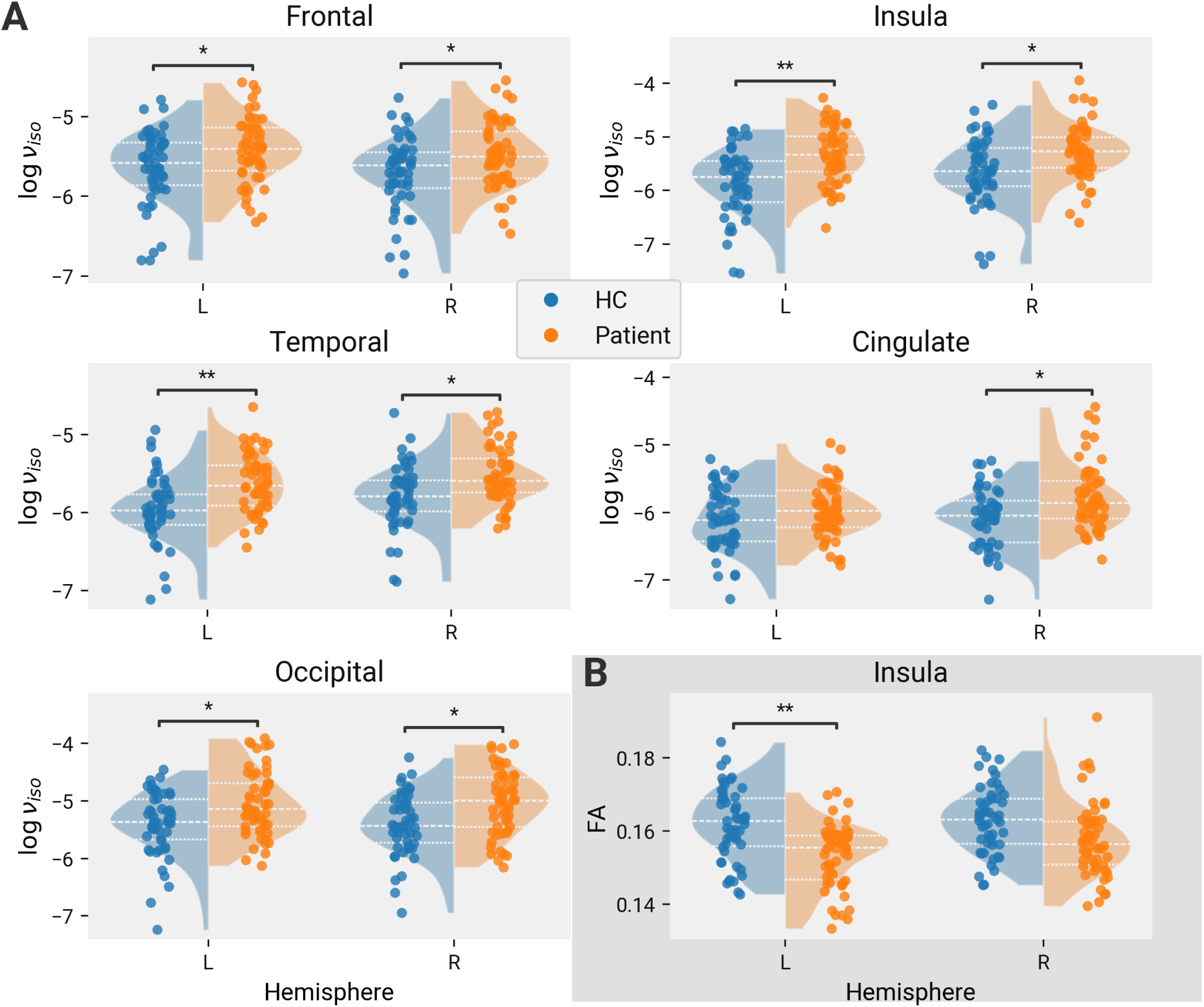
Cortical ROIs affected in early psychosis. P-values corrected for multiple comparisons across ROIs for each parameter using FDR. Left and right hemispheres are illustrated separately. For each hemisphere, data from healthy controls are shown in the left curve, patients in the right. Curves correspond to kernel density estimates with bandwidth determined using the Scott method (88). Dashed lines represent first, second, and third quartiles. A. ROIs with significantly higher *v_iso_* in both hemispheres. B. The insula has lower FA in the left hemisphere only. Statistics shown in Table 4.

**Table 4:**
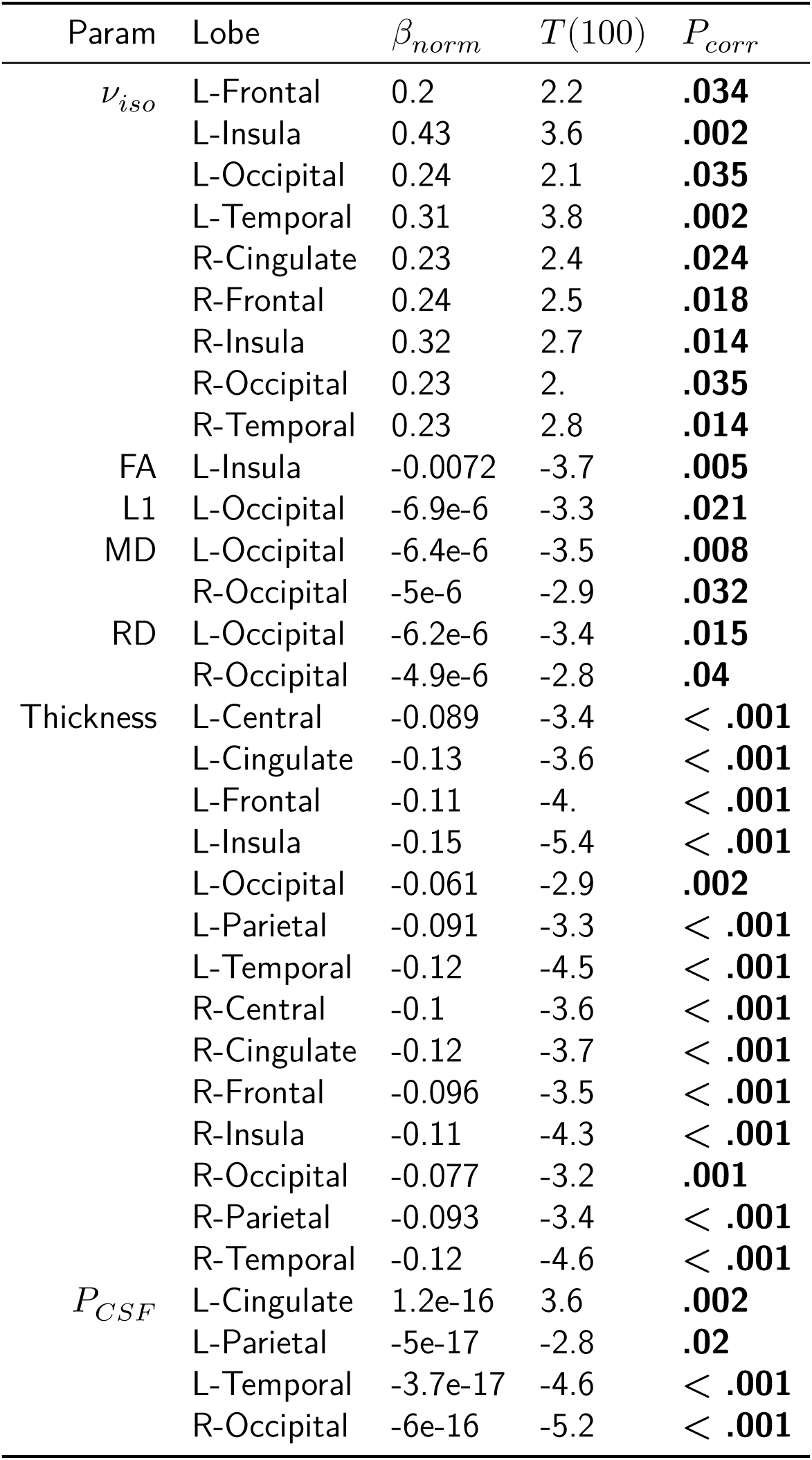
ROIs with significant group effects.

To further localize the changes, we used a random field theory (RFT) approach to identify clusters with significantly changed parameters, corrected to a FWER of 0.05. Ten clusters with significantly reduced thickness were found spanning the lobes of both hemispheres (Figure 5). We found eight clusters with increased *v_iso_* with a more restricted spatial extent, covering the left insula, posterior superior temporal gyrus, anterior inferior temporal gyrus, and occipital lobe, and the right medial orbito-frontal lobe, posterior inferior temporal gyrus, occipital lobe, and always (Figure 6).

**Figure 5:**
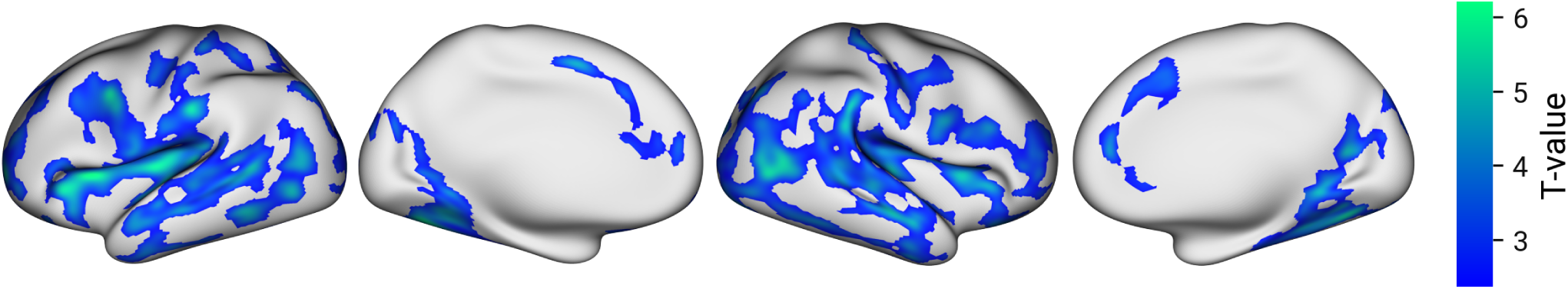
Cortical clusters with significantly lower thickness in patients. Clusters were determined with random field theory with a P-value threshold of 0.01. One-tailed t-tests were used to test for decreased thickness.

**Figure 6:**
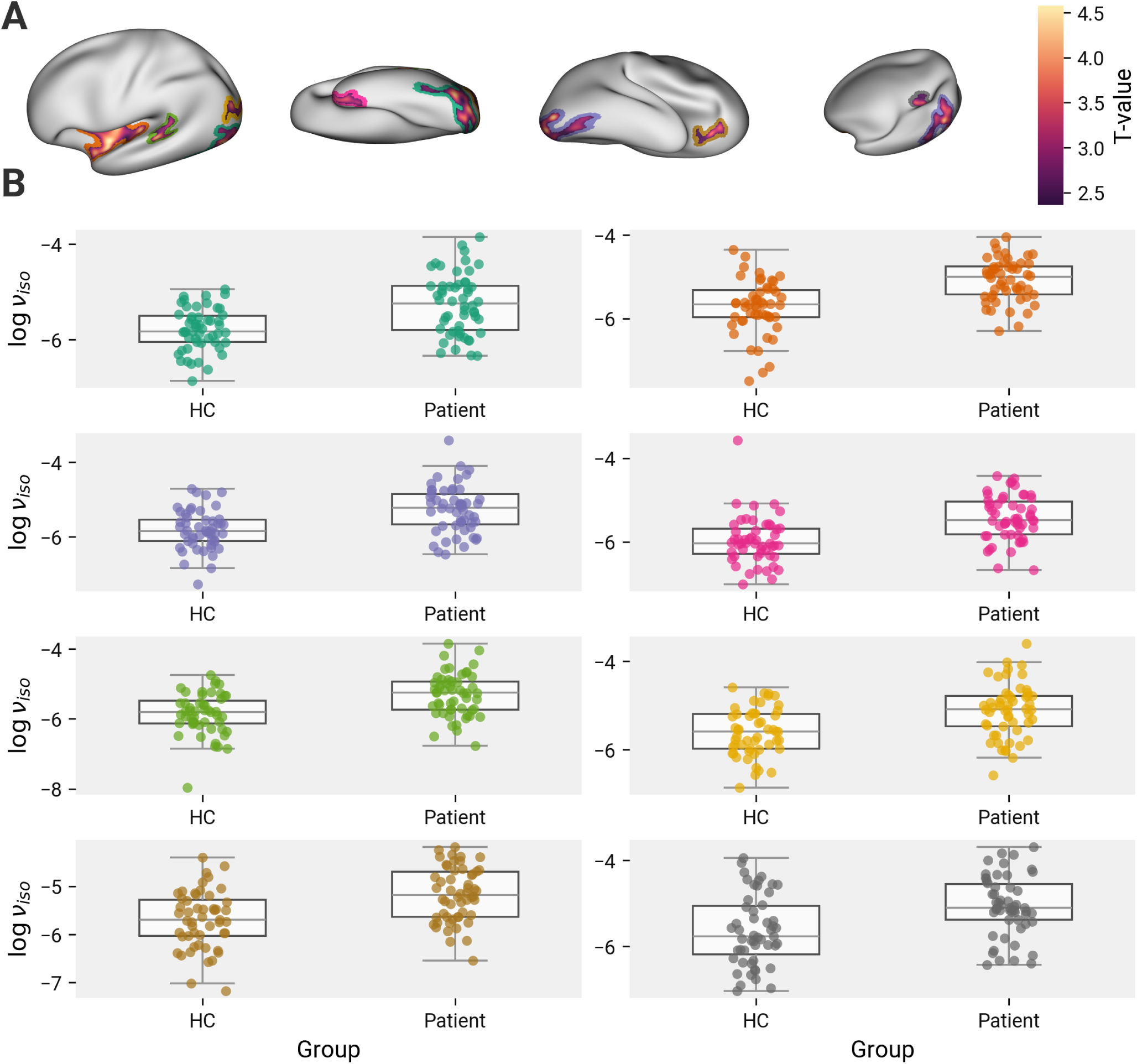
Cortical clusters with significantly higher *v_iso_* in patients. Clusters determined with random field theory with a P-value threshold of 0.01. Significance determined using one-tailed t-tests. A. Clusters with significantly higher log *v_iso_*. Colored outlines around the clusters correspond with scatter plots in B., and are not a part of the cluster. B. log *v_iso_* in individual clusters. Dots show average log *v_iso_* within clusters for individual subjects. Colors correspond with the cluster outlines in A.

Because the increase in cortical *v_iso_* was observed with a concomitant decrease in cortical thickness, there was a possibility the former was caused by increased partial volume effects in the patient group. The thinner cortical ribbon may have proportionately lower contrast with the surrounding CSF, with its very high *v_iso_*. Indeed, *v_iso_* and cortical thickness were correlated, both across subjects when comparing the subject-wise global average of the two metrics (*r*(105) = −0.57; *P* < .001) (Figure 7 A), and across the brain, using a spin-test to compare the spatial distribution of thickness and *v_fw_* across the mean parameter maps (1000 permutations; P=.028) (Figure 7 B).

**Figure 7:**
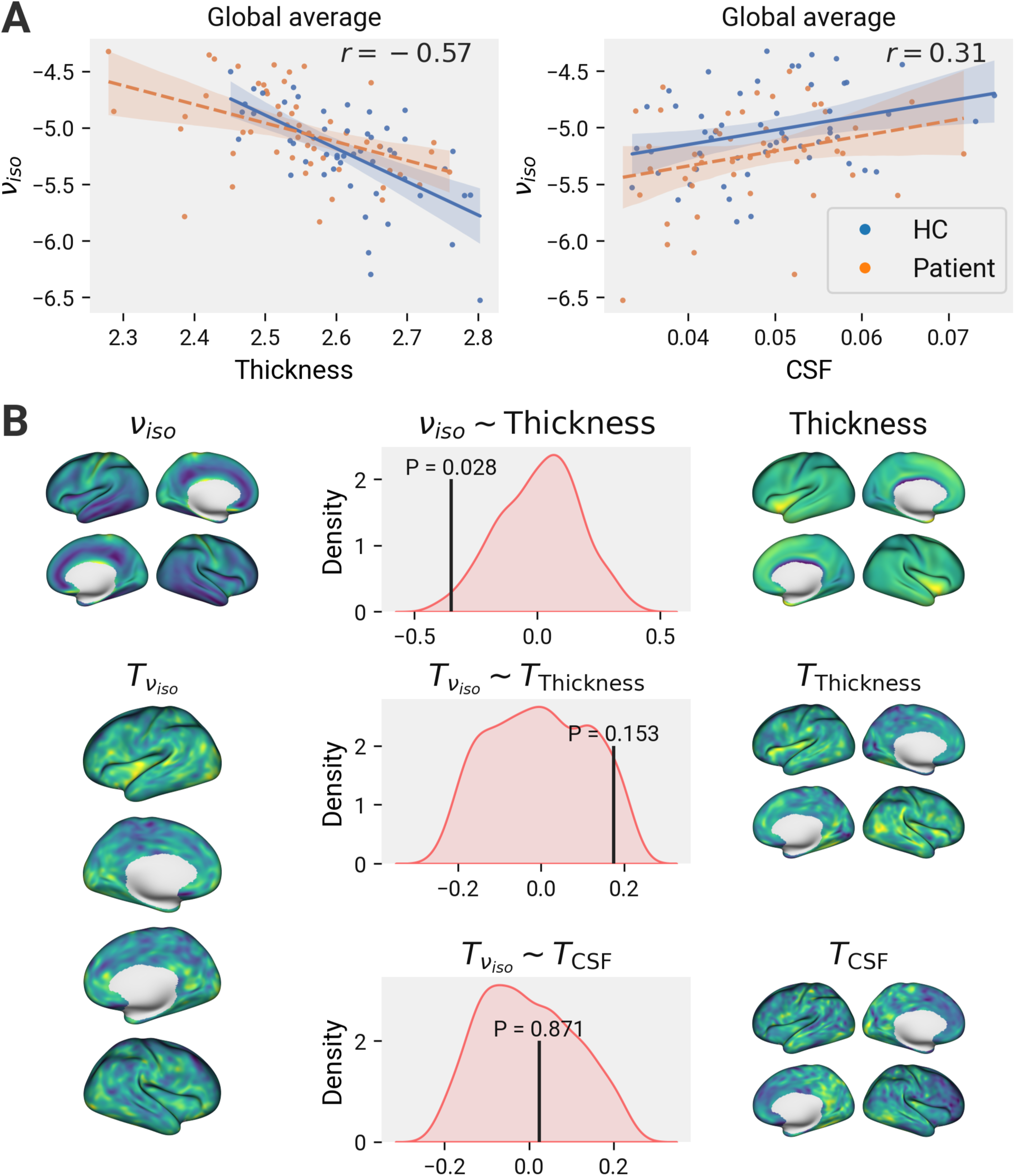
*v_iso_* findings are distinct from partial volume effects. A. Significant correlations observed across subjects between global averages of *v_iso_* and cortical thickness (*r*(105) = −0.57; *P* < .001) and CSF volume fraction (*r*(105) = 0.31; *p* = .001). Healthy controls and patients are plotted separately, but Pearson’s R is calculated over the combined sample. B. Thickness and *v_iso_* maps were made by averaging across subjects. T-value maps were made using a vertex-wise t-tests between patients and healthy controls, using age and sex as covariates. The center column shows the results of spin tests between parameter maps, testing for spatial correlation. 1000 replicates were performed per test. The correlation between parameters was tested using Spearman’s rank test. Red curves show kernel density estimate of null distribution. The empirical value is represented by the vertical line. P-value shows result of two-tailed test. The spatial distribution of thickness and *v_iso_* are significantly spatially correlated, but not the T-maps that represent diagnostic effect for *v_iso_* and thickness, or for *v_iso_* and CSF volume fraction.

We took two measures to confirm the co-occurrence of these results was not due to partial volume effects. First, we used a spin-test to compare the T-value distributions across the cortex for both the *v_iso_* and thickness group comparisons (Figure 7 B). We did not find a significant spatial correlation between the two. Second, we used *FAST* from *FSL* to generate a probabilistic CSF segmentation from the T1w image for each subject. We then sampled this CSF fraction onto cortical surface using the same procedure as the other parameters. There was no difference between groups in cortical CSF fraction either globally, in any ROI (after FDR correction), or using a RFT cluster-based approach (Figure 7 A). Finally, we compared the T-value map for the CSF fraction group comparison with the *v_iso_* map using a spin-test, and found no spatial correlation between the two (Figure 7 B).

### 3.3 White matter

To investigate changes of NODDI parameters in first episode patients, we used a composite atlas to sample NDI, ODI, and log *v_iso_* from various white matter ROIs after projecting each parameter map to a FA-derived skeleton (58). The composite atlas comprised the JHU white matter atlas, which captured core white matter regions, and the Talairach lobe segmentation, which captured peripheral white matter regions. To increase our sensitivity to local and global changes, the ROIs were organized into a hierarchical classification system. The first level was a single ROI covering the entire white matter skeleton. The second split the white matter into core and peripheral regions. The third included the peripheral lobe ROIs derived from the Talairach segmentation and four groupings of the JHU atlas labelled as projection, association, callosal, and limbic tracts. The final level included the individual JHU ROIs. Multiple comparisons were corrected separately across each level using FDR.

No changes in any of NDI, ODI, or *v_fw_* were found globally, or in any ROI at any level of the hierarchy (Figure 8). We also performed probabilistic tractography on the data, sampled the NODDI parameters underlying each streamline, and partitioned the tractogram into a connectome based on the Brainnetome atlas (51), weighted by the sampled NODDI parameters. NBS was used to test for group differences in connectome edge weights. No significant subnetworks were found for any of the parameters.

**Figure 8:**
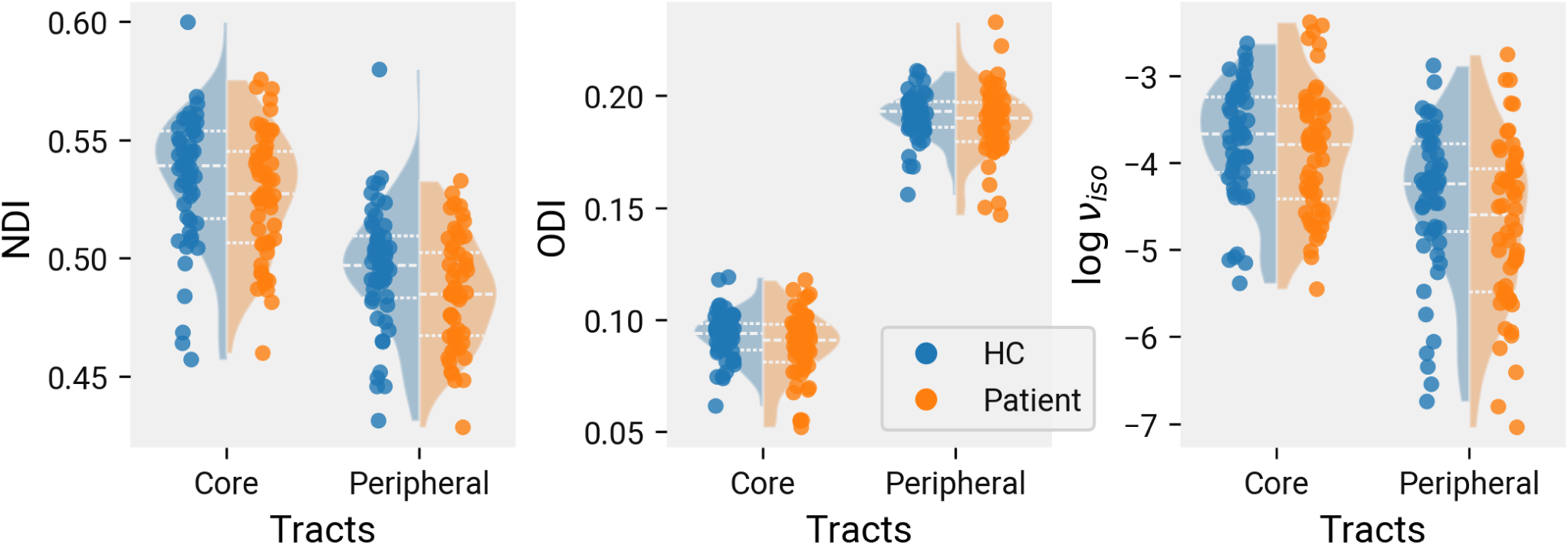
No effect of diagnosis in white matter &NODDI. Average parameter values are represented split by core and peripheral white matter. For each split, healthy controls are shown in the left curve, patients in the right. Curves correspond to kernel density estimates with bandwidth determined using the Scott method (88). Dashed lines represent first, second, and third quartiles. Diagnosis of early-stage psychosis does not significantly affect global NDI, ODI, or *v_iso_*.

### 3.4 Clinical Correlations

All of the above analysis were repeated in the patient group testing for an effect of clinical symptom scores on NODDI parameter values. We tested the PANSS30-P and PANSS30-N scores, and the disorganization factor. None of these scores had significant correlation with any NODDI parameter in the grey matter or cortical thickness, using either an ROI approach or RFT, or in the white matter, using an ROI approach or NBS.

## 4 Discussion

In our analysis of early schizophrenia patients, we observed widespread reductions in cortical thickness, in line with other reports, along with increased *v_iso_* throughout the grey matter. The prominent free water changes in the left insula coincided with a decrease in FA. As in our previous investigation of DTI metrics (1), we did not observe any differences in the white matter. We furthermore did not observe any correlations with clinical symptoms.

Our lack of findings with ODI and NDI contrasts with previous reports (21,22). Rae et al., in particular, found extensive reductions of NDI which we failed to replicate. Kraguljac et al. found more heterogeneous results. As with the DTI literature, the disparity of findings likely arises from patient heterogeneity across studies (e.g. inclusion/exclusion criteria, regional differences between patients, etc). For instance, Kraguljac et al. previously reported extensive FA reductions in the same study used in the above NODDI analysis (4). A previous pair of studies from the same authors gives the same picture. In this case, only one small cluster each of FA and ODI change were found in the same dataset (89,90). So too, in the study by (21), NDI and FA changes largely coincided. Thus, the effect size of NODDI findings may not diverge widely from FA findings within a single dataset, making the prospect of a unique sensitivity of NODDI to schizophrenia-related changes less likely.

Our NODDI findings in the grey matter of early psychosis are not as extensive as previously shown in chronic patients. We observed some increase in *v_iso_* across the entire cortex, but especially in the left insula. This finding is consistent with the synaptic density hypothesis, which predicts, either through the loss of synapses or a consequent loss of interneuron density, an increase in free water space (91). Nevertheless, the increase in *v_iso_* was spatially limited compared to thickness reduction in patients, raising the possibility of non-atrophic processes (e.g., vasogenic edema) contributing to free water increase. The particularly strong insular findings are consistent with the prominent role of the insula in schizophrenia pathophysiology (92) repeatedly highlighted by various observations, including the prominent changes in grey matter structure (26), intracortical myelin content (93), cortical folding (94) and effective connectivity (95). To our knowledge the demonstration of prominent changes in free water reflecting putative edema is a novel finding, adding an important clue that connects the suspected immunoregulatory role of the insula (96) and the emerging immune hypothesis of schizophrenia (97).

The lack of NDI and ODI changes may be explained by three considerations. First, it is not yet clear NDI would be specifically sensitive to changes in synaptic density, nor how dramatic the histological changes would need to be to drive changes on imaging. Combined NODDI and histological studies are currently scant (98), but represent an important area for future work to help interpret imaging findings. Second, it is not currently clear when a synaptic density deficit starts to appear in patients. Histological evidence comes from deceased patients well into the chronic stages of disease, and PET studies have only been done in chronic patients (91). Thus, if a synaptic density deficit only occurs later, there may no pathology to detect in early psychosis. Third, a synaptic density hypothesis does not necessarily implicate ODI, as the the elimination of neuropil is presumably omnidirectional.

Finally, we failed to find any correlations with clinical symptoms across any of the studied parameters. This contrasts with prior observations from ourselves (1) and Wang et al. (24) that increased MD and RD in the white matter correlate with higher burden of negative symptoms. Evidently, the DTI changes in patients with worse negative symptoms did not arise from fractional differences between the extra- and intra-cellular compartments, perhaps because both compartments were equally affected. The lack of correlation between symptoms and cortical thickness suggests that, while decreased thickness is a phenotype of schizophrenia as such, it does not necessarily vary with disease severity. Of course, with these conclusions come the important caveat that schizophrenia is a complex, multi-faceted disease of which the PANSS30-P and PANSS30-N scores represent only the immediate clinical aspect.

### 4.1 Limitations

Our methodology is limited by potential misalignment between the cortical surface meshes generated from the T1w anatomical scan and the diffusion scans from which we sampled our parameters. While we took efforts to ensure our results were not merely the results of partial volume effects, some level of missampling is inevitable, especially given the low thickness of the cortex relative to our diffusion scan resolution (2 mm voxels compared to thickness ranging from 1-4 mm). On the other hand, compared to previous studies which use solely voxel-based methods, our approach allowed precise inter-subject registration using the freesurfer surface-based registration approach. Furthermore, the NODDI model is only reliable to the extent its biophysical priors accurately model the underlying tissue. The extent of this accuracy is unclear: for instance, the fixed compartmental diffusivities used by the model disagree with unbounded diffusivity measurements (99).

### 4.2 Conclusion

This result confirms that extensive pre-onset disruption of white matter structure is less likely, and not necessary for the onset of psychosis. Although subtle microstructural changes (e.g. free-water excess) and the role of white matter pathology cannot be ruled out as a mediating factor in illness severity, we did not observe symptom-microstructure correlations in this study.

## Supporting information

Global Surface Statistics

ROI Surface Statistics

White Matter ROI Statistics

## Acknowledgements

Special thanks to Ravi Menon for his leadership for the imaging component of TOPSY cohort. We also thank Peter Williamson, Jean Theberge, Robert Bartha, Robert Hegele, Justin Hicks, Richard Neufeld and Keith St. Lawrence for their assistance in funding acquisition for this study.

